# SPOCK1 promotes breast cancer progression via interacting with SIX1 and activating AKT/mTOR signaling pathway

**DOI:** 10.1101/834135

**Authors:** Ming Xu, Xianglan Zhang, Songnan Zhang, Junjie Piao, Yang Yang, Xinyue Wang, Zhenhua Lin

## Abstract

SPOCK1 is highly expressed in many types of cancer, which has been recognized as a promoter of cancer progression, while its regulatory mechanism remains to be clear in breast cancer (BC). This study aimed to explore the precise function of SPOCK1 in BC progression and the mechanism by which SPOCK1 was involved in cell proliferation and epithelial-mesenchymal transition (EMT). Immunohistochemistry (IHC) and database analysis displayed that high expression of SPOCK1 was positively associated with histological grade, lymph node metastasis (LN) and poor clinical prognosis in BC. A series of assays both *in vitro* and *in vivo* elucidated that altering SPOCK1 level led to distinctly changes in BC cell proliferation and metastasis. Investigations of potential mechanisms revealed that SPOCK1 interacted with SIX1 could enhance cell proliferation, cell cycle and EMT process by activating the AKT/mTOR pathway, whereas inhibition of AKT/mTOR pathway or depletion of SIX1 reversed the effects of SPOCK1 overexpression. Furthermore, SPOCK1 and SIX1 were highly expressed in BC and might indicate poor prognoses. Altogether, SPOCK1/SIX1 promoted BC progression by activating AKT/mTOR pathway to accelerate cell proliferation and metastasis in BC, and SPOCK1/SIX1 might be promising clinical therapeutic targets to prevent BC progression.

**IMPORTANCE:** The incidence of BC is alarmingly high and many patients initially diagnosed without detectable metastases will eventually develop metastatic lesions. The occurrence of metastasis is responsible for the death of many patients, which also represents a big challenge for researchers to improve the survival rates of BC patients. Hence the scientific community pays more attention on cancer targeted therapy. This research is significant for identifying the underlying mechanisms and capabilities of SPOCK1-induced BC activities, which will greatly apply novel targets and new treatment strategies for clinicians, leading to broader biomedical impacts.

---

Breast cancer (BC), the most frequently diagnosed lethal gynecologic malignancy in women worldwide (1), which is an increasing concern because of its rising incidence. Despite major advances in BC therapy, the prognosis for most patients would be significantly worse once spread metastasized occurs (2). Metastasis is the mainspring for most patient death and represents the fundamental challenge of the clinical treatment for the patients with BC. The initiation and metastasis of BC are intricate processes, which triggered by multiple genes and intracellular signal transduction cross-talks. Thus, looking for valid molecular hallmarks and understanding the mechanisms of BC initiation and metastasis process are urged to put on the agenda.

Sparc/osteonectin, cwcv and kazal-like domains proteoglycan 1 (SPOCK1), is also known as TIC1; SPOCK; TESTICAN, belongs to multidomain testicular proteoglycan family (3). This protein family includes SPARC, TESTICAN-2, and TESTICAN-3, which are associated with cell proliferation and metastasis (4). SPARC has been properly reported in a variety of cancers (5, 6), which emphasized the involvement in cell proliferation, angiogenesis and epithelial-to-mesenchymal transition (EMT). Recent discoveries have revealed that SPOCK1 was overexpressed in colorectal cancer, non-small cell lung cancer and glioblastoma (7–9). Meanwhile, some reports revealed that SPOCK1 may promote the invasion and metastasis of gastric cancer, glioma (10, 11), and may confer a poor prognosis in urothelial carcinoma (12). Nevertheless, the underlying mechanisms and functions of SPOCK1-induced BC activities, including cancer development and metastasis process are far from clear. Here, we described the oncogenicity of SPOCK1 and clarified the molecular mechanism of SPOCK1 involved in BC evolution.

EMT was initially mentioned in embryogenesis, a kind of reversible and rapid changes cell phenotype, which is defined as the changes of the epithelial phenotype into mesenchymal features (13), including loss of contact inhibition ability, promoting cell motility and invasiveness (14). It is noteworthy that SIX homeobox 1 (SIX1), also known as BOS3; TIP39; DFNA23, an indispensable transcription factor of organogenesis (15), plays a fatal role in promoting cell EMT process (16, 17). SIX1 expression is negligible in normal adult organs, and its aberrant expression may lead to carcinogenesis (18). Recently, SIX1 was found to be referred to cellular proliferation, invasion and Warburg effect (16, 19). In BC, SIX1 was highly expressed in half of primary cancer and 90% of cancer metastasis (20). Besides, SIX1 was contributed to the initiation and prognostic of tumors (21). To date, there are no similar reports regarding the association between SIX1 and SPOCK1 in BC evolution.

Herein, we aimed to reveal that overexpression of SPOCK1/SIX1 was related to BC cell proliferation and metastasis and predicted poor prognosis in BC patients via bioinformatic analysis of available BC datasets and immunohistochemical (IHC) assay. Additionally, we demonstrated that SPOCK1/SIX1 activated PI3K/AKT/mTOR pathway, consequently promoting BC cell proliferation, accelerating cell cycle progression, trigger cell EMT program and metastasis, which provided a new strategy for BC targeted therapy.

## RESULTS

### SPOCK1 was abnormally and strongly expressed and associated with metastasis and poor prognostic in BC

SPOCK1 was highly expressed in various cancers, prostate cancer, pancreatic cancer, lung cancer and breast cancer included (Fig. 1A). We analyzed SPOCK1 mRNA expression across different datasets (27–29) on Oncomine database and found that SPOCK1 was higher-expressed in BC than in normal tissues (Fig. 1B). The median rank of SPOCK1 in high expressed genes of BC was 672.0 based on a meta-analysis across seven datasets for Oncomine algorithms (*P*=1.06E-11) (Fig. 1C). Furthermore, the UALCAN database, which integrated TCGA samples, showed the same results (Fig. 1D). UALCAN also displayed mRNA expression of SPOCK1 in 33 kinds of cancers, in which several cancers increased expressed SPOCK1, including breast cancer. In the HPA database, SPOCK1 protein expression was manifested hardly detected in normal sections but significantly higher expressed in BC (Fig. 1E).

**Fig. 1.**
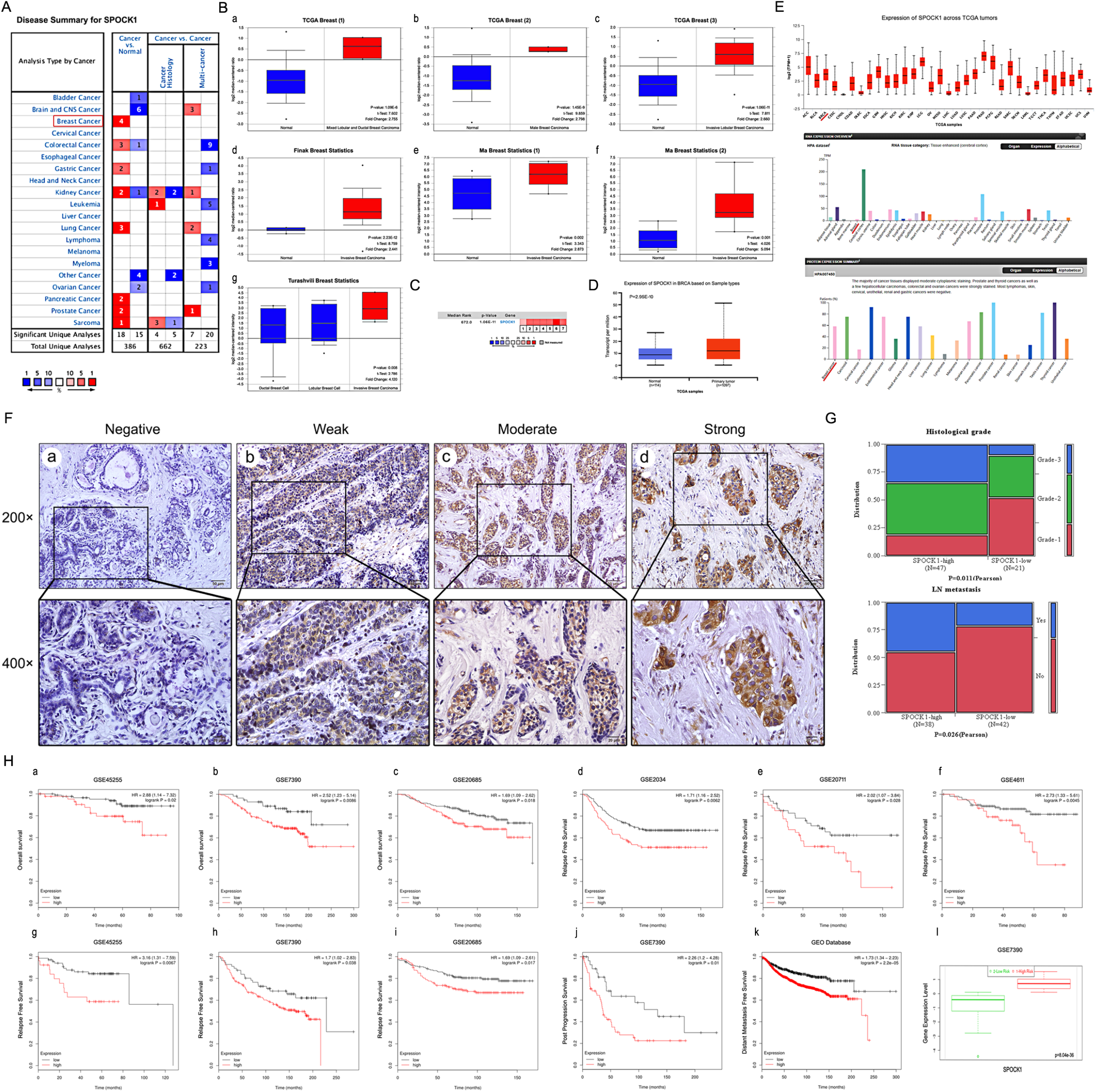
Overexpression of SPOCK1 is positively associated with histological grade, LN metastasis and poor prognosis in BC. **A:** The graphic showed the numbers of datasets with statistically significant mRNA high expression (red) or down-expression (blue) of SPOCK1 (cancer vs. Normal tissue). The *P*-value threshold was 0.01. **B:** Box plots derived from gene expression data in Oncomine comparing expression of SPOCK1 in normal and BC tissue. The *P*-value was set up at 0.01 and fold change was defined as 2. **C:** A meta-analysis of SPOCK1 gene expression from seven Oncomine databases where colored squares indicated the median rank for SPOCK1 (*vs*. Normal tissue) across 7 analyses. **D:** The expression of SPOCK1 was elevated in BC compared to normal breast tissues. Data derived from UALCAN database. **E:** Expression of SPOCK1 across TCGA carcinomas from Ualcan database (a); overview of SPOCK1 protein levels in BC tissues and normal breast tissues (b-c). **F:** IHC staining (negative, weak, moderate and strong expression) for SPOCK1 in BC tissues (a-d). **G:** Relationships between SPOCK1 expression and clinicopathologically significant aspects of BC. **H:** Overall survival (OS) (a-c), relapse free survival (RFS) (d-i), post progression survival (PPS) (j), distant metastasis free survival (DMFS) (k) and risk assessment curves (l) of patients with or without elevated SPOCK1 levels. Survival data derived from Kaplan–Meier (KM) plotter database. High SPOCK1 expression levels were found in high risk groups of BC patients. Box plots generated by SurvExpress showed the expression levels of SPOCK1 in indicated dataset and the *P*-value resulting from a *t*-test. Low-risk groups are denoted in green and high-risk groups in red, respectively.

To further confirm the expression pattern of SPOCK1 in BC, 80 BC tissues and 10 adjacent non-tumor tissues were examined by IHC assay. IHC analysis showed that SPOCK1 was significantly higher expressed in BC tissues than in adjacent non-tumor ones. The positive rate (93.8%; 75/80) and strongly positive rate (72.5%; 58/80) of SPOCK1 in BC were both significantly higher than in adjacent non-tumor tissues (30.0%, 3/10 and 10%; 1/10) (*P*<0.001) (Fig. 1F, Table 1), which confirmed that SPOCK1 was aberrantly upregulated in BC. Notably, aberrant SPOCK1 expression was associated with histological differentiation (*P*=0.011), and LN metastasis (*P*=0.026), but not with the patients’ age or the status of ER and PR (Fig. 1G, Table 2). Not only that, but high SPOCK1 expression was markedly related to unfavorable outcomes in BC patients. We evaluated the relationship between the SPOCK1 expression level and OS, RFS, PPS and DMFS of patients with BC by Kaplan-Meier plotter database. As showed in Fig. 1H, high SPOCK1 expression resulted in shorter OS, RFS, PPS and DMFS in various datasets. Finally, the level of SPOCK1 was significantly higher in the high-risk group compared to low via SurvExpress database. In general, these results underscored that SPOCK1 was strongly expressed in BC and could serve as an outcome predictor in BCs.

**Table 1.** SPOCK1 expression in BC

| Diagnosis | No. of case | Positive cases |  |  |  | Positive rates | Strongly positive rates |
| --- | --- | --- | --- | --- | --- | --- | --- |
|  |  | - | + | ++ | +++ |  |  |
| **Breast cancer** | 80 | 5 | 17 | 26 | 32 | 93.8% | 72.5% |
| **Adjacent non-tumor** | 10 | 4 | 2 | 1 | 0 | 30.0% | 10.0% |

\* $P < 0.05$ and \*\* $P < 0.01$ : compared with adjacent non-tumor

**Table 2.** Relationship between SPOCK1 expression and clinicopathologic features of BC patients

| Variables | No. of case | SPOCK1 strongly positive cases (%) | $\chi^2$ | $P$ value |
| --- | --- | --- | --- | --- |
| **Age** |  |  |  |  |
| ≥52 | 43 | 74.4%(32/43) | 0.172 | 0.679 |
| <52 | 37 | 70.3%(26/37) |  |  |
| **Histological grade** |  |  |  |  |
| Grade-1 | 20 | 45%(9/20) | 8.996 | 0.011* |
| Grade-2 | 30 | 73.3%(22/30) |  |  |
| Grade-3 | 18 | 88.9%(16/18) |  |  |
| **ER** |  |  |  |  |
| Positive | 50 | 70%(35/50) | 0.418 | 0.518 |
| Negative | 30 | 76.7%(23/30) |  |  |
| **PR** |  |  |  |  |
| Positive | 54 | 70.4%(38/54) | 0.653 | 0.419 |
| Negative | 24 | 79.2%(19/24) |  |  |
| **LN metastasis** |  |  |  |  |
| Yes | 26 | 65.4%(17/26) | 4.941 | 0.026* |
| No | 54 | 38.9%(21/54) |  |  |

\* $P < 0.05$ and \*\* $P < 0.01$

### SPOCK1 accelerated cell cycle progression and promoted cell proliferation in BC

To verify the potential oncogenic activity of SPOCK1 in BC, we surveyed SPOCK1 endogenous expression in a series of BC cell lines and a normal immortalized mammary gland cell line by western blot. Among 7 cell lines, MCF7 and SKBR3 cell lines exhibited high SPOCK1 expression, MDA-MB-231 and HS 578T cell lines showed low expression (Fig. 2A). To explore the potential biologic function of SPOCK1, we chose MCF7 and SKBR3 cell lines for SPOCK1 knockdown, chose MDA-MB-231 and HS 578T cell lines for SPOCK1 stable overexpression. The level of SPOCK1 expression in stable infected cell lines was verified by western blot (Fig. 2B), and the transfection efficiency was shown in Fig. 2C. The best silencing effect was obtained with the shSPOCK1#2 and shSPOCK1#3 constructions for MCF7 and SKBR3 cell lines. Meanwhile, stability overexpression of SPOCK1 in MDA-MB-231 and HS 578T cell was exhibited.

**Fig. 2.**
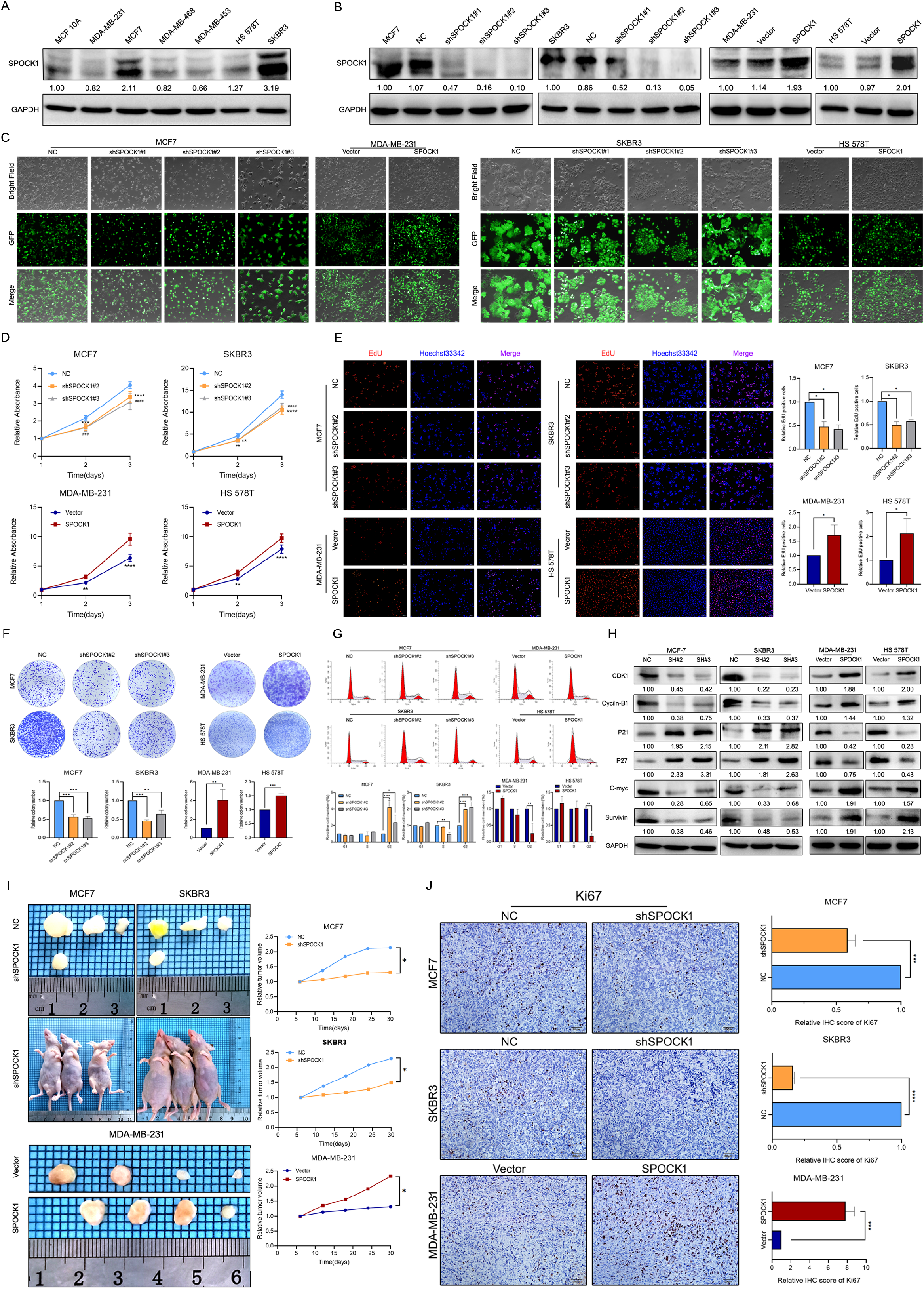
SPOCK1 influences BC cell growth. **A:** Protein expression levels of SPOCK1 in BC cell lines as determined by western blot analysis. **B:** MCF7/SKBR3 cells with SPOCK1 silencing and MDA-MB-231/HS 578T cells with SPOCK1 overexpression were established by viral transduction. The SPOCK1 levels in these established cell lines were verified by western blot analysis at 48 h after transfection. **C:** Cells in bright light and GFP were captured to merge for displaying the transfection efficiency. **D:** Cell viability was examined by MTT assay. **E:** Results of EdU assay on BC cells. Representative photographs are shown at the original magnification, ×100. **F:** Cell clonogenic capacity was measured by colony formation assay. **G:** Flow-cytometry analysis was performed to detect cell cycle progression. **H:** The expression of proteins related cell cycle (CDK1, Cyclin-B1, P21, P27, C-mvc and Survivin) was determined by western blot analysis. GAPDH was used as a loading control. **I:** Xenograft tumors formed by injecting the indicated cells. Relative tumor volume curves were summarized in the line chart (\**P*<0.05). **J:** IHC staining of the proliferation marker Ki67 in xenograft tumors. The relative percentage of Ki67-positive cells was summarized in the bar charts. The P values were obtained using t-tests (\*\*\**P*<0.001, \*\*\*\**P*<0.0001). All results are from three independent experiments. The error bars represent the SD.

MTT and EdU incorporation assays were used to determine the potential function of SPOCK1 in BC proliferation. As showed in Fig. 2D-E, silencing SPOCK1 significantly suppressed the cell growth, whereas SPOCK1 overexpression promoted cell proliferation. Similarly, upregulation of SPOCK1 facilitated the cell clonogenicity, while SPOCK1 knockdown formed the smaller and fewer colonies compared with the controls (Fig. 2F). We further analyzed the influence of cell cycle with different levels of SPOCK1 expression. The flow cytometry assay indicated that cells with higher SPOCK1 expression accelerated the progress of G_2_/M phase, compared with respective control (Fig. 2G). Additionally, SPOCK1 knockdown decreased the level of Cyclin-B1, CDK1, C-myc and Survivin, which were coupled with a concomitant increasing the expression of P21 and P27 (Fig. 2H). Conversely, ectopic expression of SPOCK1 displayed adverse results. Together, the results demonstrated that SPOCK1 plays a crucial role in BC cell cycle and proliferation *in vitro*.

*In vivo*, stable BC cells with modified SPOCK1 expression were subcutaneously injected into the fourth mammary fat pad of nude mice. The volumes of tumors formed by high SPOCK1 expression cells were significantly higher than those from low expression cells (Fig. 2I). Additionally, the expression of Ki67 in MCF7-NC group SKBR3-NC group and MDA-MB-231-SPOCK1 group was much higher than in their negative control groups (Fig. 2J). Overall, these findings suggested that SPOCK1 promoted the BC cell growth *in vivo*.

### SPOCK1 motivated BC metastasis via EMT process

Next, we observed the metastatic ability of BC cells with different SPOCK1 expression. The wound-healing assay indicated that cells with higher SPOCK1 expression displayed a more widespread wound closure area, compared with corresponding control (Fig. 3A). Transwell assays provided consistent evidence (Fig. 3B). To further explore the effect of SPOCK1 on BC metastasis *in vivo*, stable cells with modified SPOCK1 expression were injected into nude mice tail vein. The number of pulmonary metastasis in higher SPOCK1 expression group was significantly more than corresponding control, which got the opposite result in lower SPOCK1 expression group (Fig. 3C). Meanwhile we found that the cells with higher SPOCK1 expression lost cell polarity, displayed spindle-shape and acquired mesenchymal morphology of stronger invasion and metastasis ability but lower SPOCK1 expression tended to be an opposite morphology (Fig. 3D).

**Figure 3:**
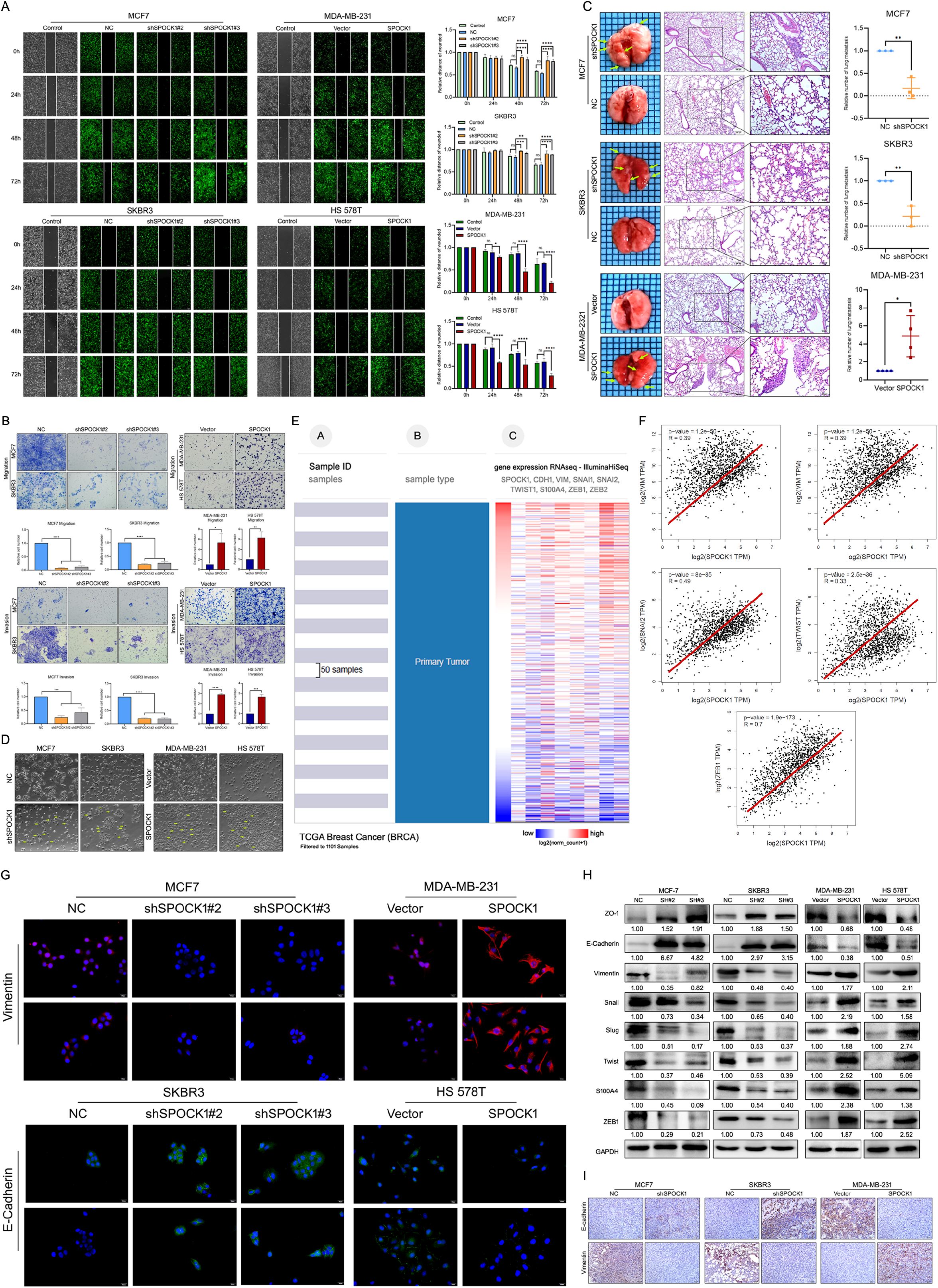
SPOCK1 promotes cellular invasion, metastasis and the EMT *in vitro*. **A:** A scratch wound-healing assay was used to determine the effects of SPOCK1 on BC cell motility. **B:** Results of a transwell migration assay (a) and a Matrigel invasion assay (b) for cellular invasion. The mean number of cells in five fields per membrane is shown (×200). **C:** Representative images of gross and hematoxylin and eosin (H&E) staining and relative numbers of lung surface metastatic foci detected in each group (*P<0.5, **P<0.01). The scale bar is 100 μM and 50 μM. **D:** Representative images showing the morphological changes in the indicated cell lines. **E:** The heat maps of the correlation between SPOCK1 and EMT markers in the same cohort. **F:** Positive relationships for SPOCK1 and EMT markers were showed on GEPIA2. **G:** The expression of EMT markers was detected by immunofluorescence staining in BC cells. The scale bar is 20 μM. **H:** The expression of epithelial markers (E-cadherin and ZO-1) and mesenchymal markers (Vimentin, Snail, Slug, Twist, S100A4 and ZEB1) was determined by western blot analysis. GAPDH was used as a loading control. **I:** IHC staining for E-cadherin and Vimentin protein in tumor specimens from xenografts (200×). The P values were obtained using Mann-Whitney U tests or t-tests (\**P*<0.05, \*\**P*<0.01, \*\*\**P*<0.001, \*\*\*\**P*<0.0001). All results are from three independent experiments. The error bars represent the SD.

To speculate whether EMT should be responsible for SPOCK1-mediated changes in BC metastasis, we analyzed a cohort of 1101 BC samples from TCGA dataset by UCSC Cancer Genomics Browser. As presented in Fig. 3E, we found that the heat maps of SPOCK1 in BC were strikingly coincident with VIM, SNAI1, SNAI2, TWIST1, S100A4, ZEB1, and ZEB2, inversely proportional to CDH1 (E-cadherin). Similarly, GEPIA2 database also showed positive correlations between SPOCK1 and mesenchymal proteins (Fig. 3F). Then western blot and IF assays showed that downregulation of SPOCK1 accelerated the expression of epithelial markers and accompanied by a reduction of mesenchymal markers (Fig. 3G-H). Consistently, the group with high SPOCK1 expression displayed inverse results. Additionally, IHC staining results showed a higher expression of E-Cadherin and lower expression of Vimentin in shSPOCK1 group tumor tissue. Conversely, sections with high SPOCK1 expression displayed the opposite effects (Fig. 3I). Taken together, these findings indicated that SPOCK1 enhanced EMT progression and triggered BC metastasis *in vitro* and *in vivo*.

### The oncogenic activity of SPOCK1 was significantly correlated with AKT/mTOR pathway

AKT/mTOR signaling pathway has vital roles in cancers evolution, which activation has been found in most BCs (30–33). Thus, we speculated that whether SPOCK1 involved the regulation of AKT/mTOR pathway in BC. Strikingly, depletion of SPOCK1 resulted in a decreased abundance of p-AKT, p-mTOR, p-S6 and p-4EBP1, where the total protein levels were not influenced (Fig. 4A). Then we further explored which role did the AKT/mTOR pathway played in SPOCK1-mediated regulation of BC. We used PI3K/AKT inhibitor LY 290042 and mTOR inhibitor Rapamycin to block PI3K/AKT/mTOR activity, and found that the inhibitors not only suppressed the activation of PI3K/AKT/mTOR pathway, but also reversed the promotion of SPOCK1 to the pathway. However, the inhibitors had no effects on SPOCK1 expression (Fig. 4B). Indeed, LY 290042 and Rapamycin significantly suppressed the ability of SPOCK1 to accelerate the BC cell proliferation and cell cycle progress (Fig. 4C-G). Similarly, inhibitors almost abolished the promotion of SPOCK1 on BC cell migration, invasion and EMT progress (Fig. 4H-J). These results demonstrated that SPOCK1, at least partly, contributed to BC proliferation and EMT by activating AKT/mTOR signaling pathway.

**Figure 4:**
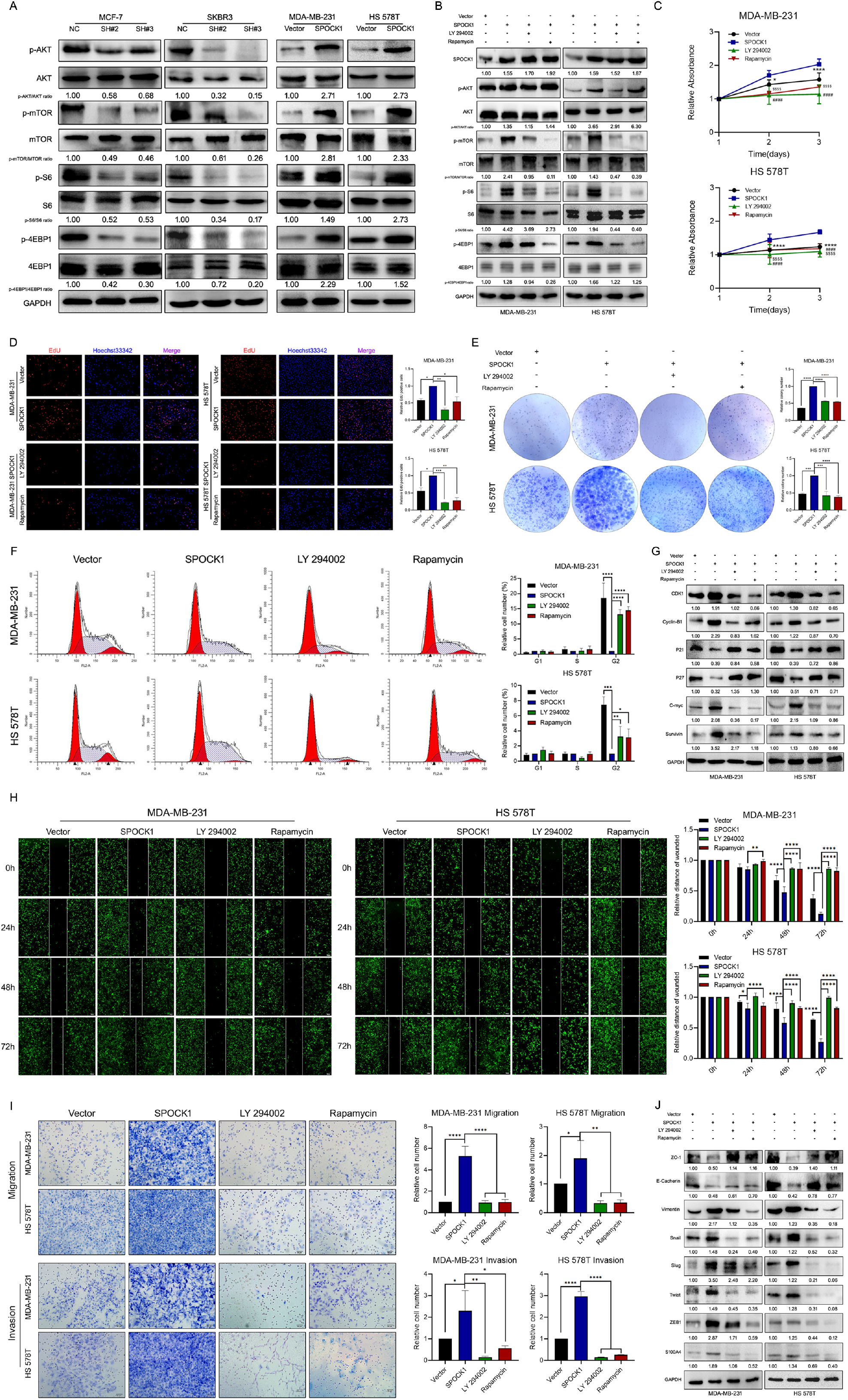
SPOCK1 activates the AKT/mTOR signaling pathway in BC cells. **A:** Proteins level on AKT/mTOR pathway of indicated cells were assayed by western blotting. GAPDH was used as a loading control. **B:** Stable BC cells were treated with LY 290042 or Rapamycin. Then indicated protein levels were assayed by western blotting. GAPDH was used as a loading control. **C-E:** Cell viability was detected in SPOCK1-overexpressed cells after treatment with LY 290042 or Rapamycin by MTT assay (C), Edu staining (D) and colony formation (E) assays. **F:** Cell cycle progression was assayed by flow-cytometry analysis after dealing with LY 290042 or Rapamycin. **G:** Stable BC cells were treated with LY 290042 or Rapamycin. Then cell cycle related protein levels were assayed by western blotting. GAPDH was used as a loading control. **H-I:** Cell motility and invasion capacities was detected in SPOCK1-overexpressed cells after treatment with rapamycin or LY294002. **J:** Stable BC cells were treated with LY 290042 or Rapamycin. Then levels of EMT-related proteins were assayed by western blotting. GAPDH was used as a loading control. (\**P*<0.05, \*\**P*<0.01, \*\*\**P*<0.001, \*\*\*\**P*<0.0001).

### Identification of SIX1 served as a target gene of SPOCK1 in BC

To further explore the molecular mechanisms underlying SPOCK1-induced BC proliferation and metastasis process, we found the potential target gene of SPOCK1 by bioinformatics strategies. We detected SPOCK1 protein-protein interactions by web-based databases, STRING and GeneMANIA, and intriguingly found that SPOCK1 was also associated with SIX1 (Fig. 5A). The mechanism has been clarified that SIX1 could induce BC cells to undergo EMT progress and metastasis via TGF-β pathway (34, 35). Moreover, SIX1 was highly expressed in breast cancer (27, 29) (Fig. 5B-E). The UALCAN and HPA databases showed the same results (Fig. 5F-G). Additionally, Kaplan Meier-plotter and SurvExpress databases displayed that high level of SIX1 expression showed poor OS, RFS and DMFS, and acquired higher risk (Fig. 5H). In all, these data highlighted that SIX1 was highly expressed in BC and correlated with poor clinical outcome.

**Figure 5:**
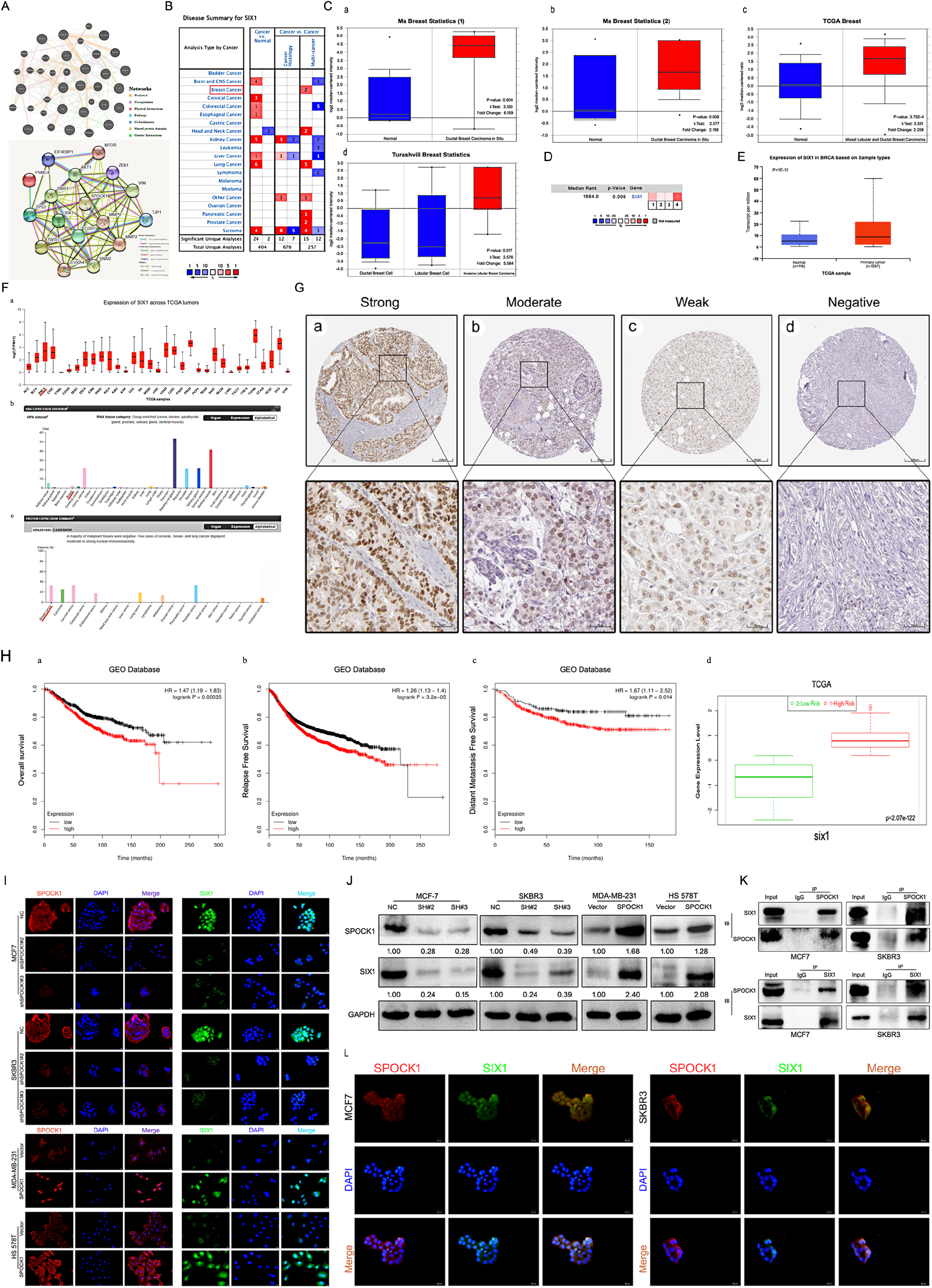
Identification of SIX1 as a downstream mediator of SPOCK1 in BC cells. **A:** Network diagram of SPOCK1/SIX1 protein interaction by GeneMANIA (a) and STRING (b). **B:** The graphic showed the numbers of datasets with statistically significant mRNA high expression (red) or down-expression (blue) of SIX1 (cancer vs. Normal tissue). The *P*-value threshold was 0.01. **C:** Box plots derived from gene expression data in Oncomine comparing expression of SIX1 in normal and BC tissue. The p value was set up at 0.01 and fold change was defined as 2. **D:** A meta-analysis of SIX1 gene expression from four Oncomine databases where colored squares indicated the median rank for SIX1 (*vs*. Normal tissue) across 4 analyses. **E:** The expression of SIX1 was elevated in BC compared to normal breast tissues. Data derived from UALCAN database. **F:** Expression of SIX1 across TCGA carcinomas from Ualcan database (a); overview of SIX1 protein levels in BC tissues and normal breast tissues (b-c). **G:** IHC staining (negative, weak, moderate and strong expression) for SIX1 in BC tissues (a-d). Data derived from HPA database. **H:** Overall survival (OS) (a), relapse free survival (RFS) (b) and distant metastasis free survival (DMFS) (c) curves of patients with or without elevated SPOCK1 levels. Data derived from Kaplan–Meier (KM) plotter database. High SPOCK1 expression levels were found in high risk groups of BC patients (d). Data derived from SurvExpress database. **I-J:** Expression levels of indicating cells were assayed by IF and western blotting. GAPDH was used as an internal control. **K:** The interaction between endogenous SPOCK1 and SIX1 proteins was analyzed by coimmunoprecipitation in MCF7 and SKBR3 cells. **L:** Immunofluorescence double-labeling experiments confirmed the existence of SPOCK1-SIX1 colocalization phenomena in the cytoplasm. The scale bar is 20 μM.

According to these data, we conjectured whether SPOCK1 enrichment in cells had an affiliation with SIX1. Consistent with our conjecture, IF and western blots assays displayed that SIX1 was enriched in SPOCK1 highly expressed cells, whereas SPOCK1 knockdown decreased the level of SIX1 expression (Fig. 5I-J). To further explore potential binding interaction between SPOCK1 and SIX1, co-immunoprecipitation (Co-IP) assay was performed. As showed in Fig. 5K, we discovered the physical interaction of SPOCK1 with SIX1. Additionally, IF staining showed colocalization of SPOCK1 and SIX1 in cytoplasm (Fig. 5L). Taken together, our findings indicated that SIX1 served as a target gene of SPOCK1 in BC.

### The SPOCK1/SIX1 axis regulated BC proliferation and metastasis via AKT/mTOR signaling activity

To explore the potential mechanism of SIX1 involved SPOCK1-induced proliferation, EMT and metastasis, we knocked-down SIX1 expression by siRNA in SPOCK1 stable overexpression cells (Fig. 6A). As expect, silencing the expression of SIX1 effectively restrained the SPOCK1-mediated cell proliferation, clone formation and cell cycle progression (Fig. 6B-E), and reversed SPOCK1-induced cell motility, migration and invasion (Fig. 6F-H). Furthermore, down-regulated the expression of SIX1 was substantially offset the SPOCK1-involved activation of AKT/mTOR signaling but didn’t affect the level of SPOCK1 expression (Fig. 6I). Altogether, this evidence suggested that SIX1 silencing could, at least partially, abolish the biological behaviors that SPOCK1-induced in BC.

**Figure 6:**
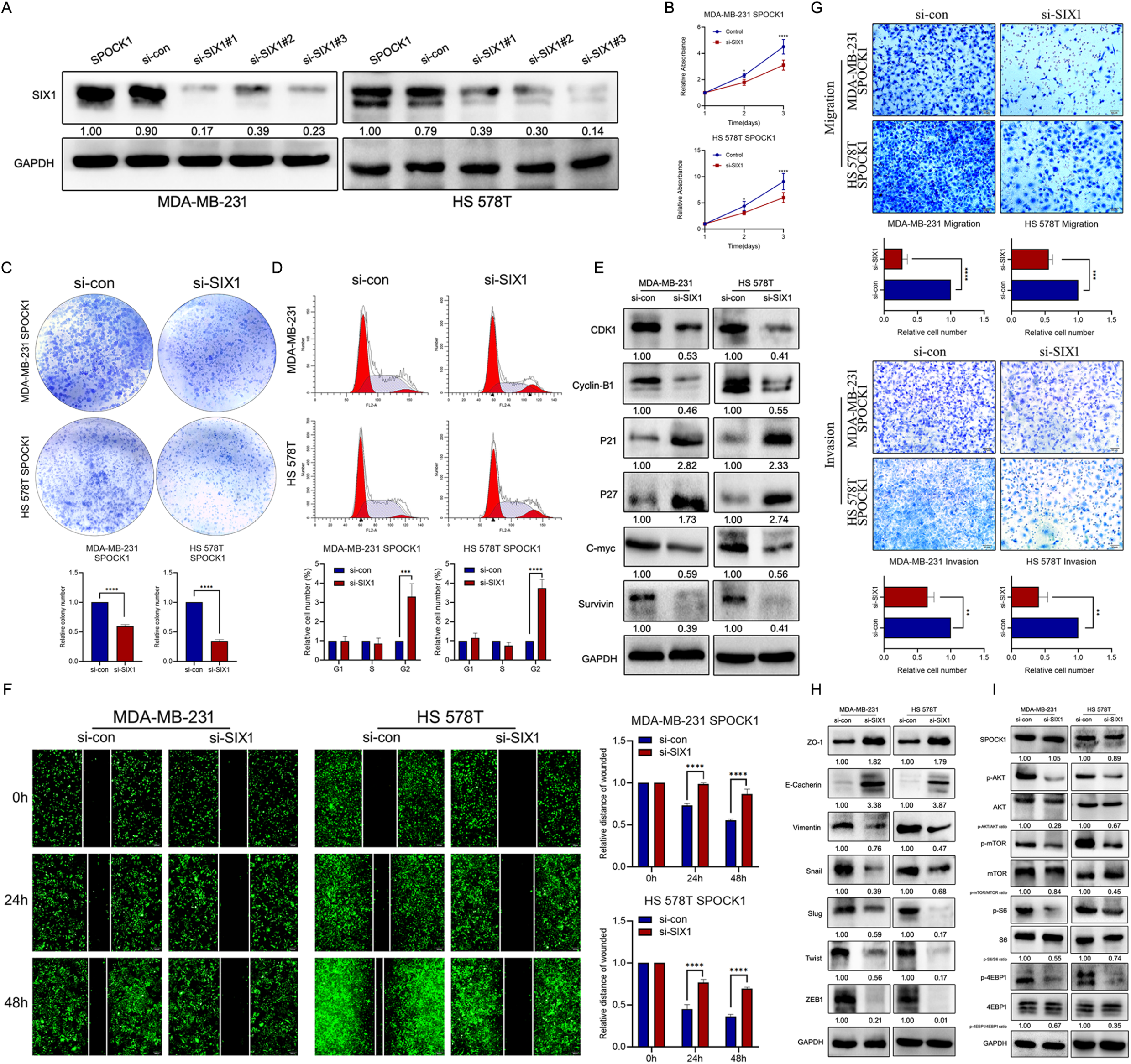
SIX1 involved in SPOCK1-mediated BC progression. **A:** MDA-MB-231 and HS 578T cell line were transduced with si-con, si-SIX1#1, si-SIX1#2 and si-SIX1#3. The SIX1 levels in these were verified by western blot analysis after 48 h transfection. **B-C:** Cell viability was detected in SPOCK1-overexpressed cells after transduction with si-RNAs by MTT assay (B) and colony formation (C) assay. **D:** Cell cycle progression was assayed by flow-cytometry analysis after dealing with si-RNAs. **E:** Stable BC cells were treated with si-RNAs. Then cell cycle related protein levels were assayed by western blotting. GAPDH was used as a loading control. **F-G:** Cell motility and invasion capacities was detected in SPOCK1-overexpressed cells after treatment with si-RNAs. **H-I:** Stable BC cells were treated with si-RNAs. Then levels of EMT-related proteins and AKT/mTOR pathway were assayed by western blotting, respectively. GAPDH was used as a loading control. (\**P*<0.05, \*\**P*<0.01, \*\*\**P*<0.001, \*\*\*\**P*<0.0001).

## DISCUSSION

SPOCK1 is an extracellular matrix glycoprotein, which is regarded as a candidate oncogene, has been verified to be closely related to the tumorigenesis, tumor progression, adhesion and metastasis of various tumors (36, 37). Chen *et al*. reported that SPOCK1 was involved in slug-induced EMT and promoted cell invasion and metastasis, which high expression was also a prognostic factor for poor survival in gastric cancer (10). Results in this study documented that SPOCK1 was highly expressed in BC cells and clinical specimens relative to normal ones, which were paralleled with Oncomine, HPA and Ualcan databases. IHC analysis displayed that SPOCK1 was related to tumor histological differentiation and LN metastasis. Moreover, Kaplan Meier-plotter database displayed that BC patients with high SPOCK1 expression had poor OS, RFS, PPS and DMFS. Our findings were similar with the results in urothelial carcinoma (UC), that higher SPOCK1 expression was correlated with unfavorable clinicopathological parameters and conferred a poor prognosis in UC (12). Evidence from SurvExpress database further showed that high expression of SPOCK1 had higher risk in BC. These observations illustrated that SPOCK1 might play a crucial role in the treatment and prognosis evaluating of BC.

Infinite proliferation and metastasis are leading responsible for tumor malignant phenotypes, while initiation of EMT is the early step of the metastatic cascade. Unquestionably, EMT is identified as the dominant program in BC initiation and metastatic spread (38, 39). A previous study revealed that deregulating the expression of SPOCK1 suppressed colorectal cancer (CRC) proliferation *in vitro* and *vivo*, SPOCK1 was involved in CRC malignant features (7). Notably, SPOCK1 was altered by EPCR to mediate 3D growth consequently promoted breast cancer progression (40). Here, we detected the variation on BC cells by modifying SPOCK1 expression, which revealed that SPOCK1 overexpression improved the proliferative and metastatic properties of BC cells and suppression of SPOCK1 got the opposite effect. The xenograft and lung metastasis models further confirmed the results *in vitro*. Specifically, western blot and IF analyses displayed that SPOCK1 gained the expression of mesenchymal markers and lost the epithelial markers. Moreover, similar evidence was verified in xenograft mouse sections, suggesting that SPOCK1 triggered EMT process both *in vitro* and *in vivo*.

The synthesis of multifarious signaling molecular events leaded the oncogenesis of BC. Understanding and identifying these signaling mechanisms would enable restrain EMT progress to further therapeutic control cancer metastasis (41–43). Previous studies reported that SPOCK1 could mediate EMT by Wnt/β-catenin signaling in non-small cell lung cancer (8), PI3K/AKT signaling pathway in colorectal cancer and others (7). Moreover, SPOCK1 blocked gallbladder cancer (GBC) cell apoptosis and promoted cell proliferation and metastasis via activating PI3K/Akt signaling both *in vitro* and *in vivo* (44). At present, therapy of BC via targeting PI3K/AKT/mTOR pathway is still in evolving field (45). Thus, we were particularly interested in exploring the special role of SPOCK1 in the evolution of normal mammary gland to BC induced by the AKT/mTOR pathway. Our study revealed that SPOCK1 overexpression actives the AKT/mTOR pathway to promote the progression of BC. This was confirmed by the alterations of target proteins of AKT/mTOR pathway induced by depleting or overexpressing SPOCK1 expression, which can be reversed by LY 290042 or Rapamycin, respectively. Furthermore, inhibition of AKT/mTOR pathway by LY 290042 or Rapamycin also impaired the effect of SPOCK1 upregulation on cell cycle, proliferation and EMT process, verifying the effect of AKT/mTOR pathway on SPOCK1-induced BC cell growth and metastasis.

As is well known that SIX1 made a notable contribution to tumor growth and metastasis (46, 47). SIX1 was identified to involve in the oncogenic role of SPOCK1 in BC. Hyperactivation of SIX1 was widespread in a variety of human tumors and associated with poor clinical efficacy. Emerging evidence revealed that SIX1 targeting the ERK and AKT signaling and promoting malignant behavior of cancer cells (16, 48). In accordance with this, we searched the online databases and observed that SIX1 was frequently high expressed in many cancers, BC included, and correlated with poor survival and high risk. Moreover, statistically interaction between SPOCK1 and SIX1 by an online gene expression profiling interactive analysis tool. Based on these findings, we paid our attention to the relationship of SPOCK1 and SIX1. As modified SPOCK1 expression led to significantly changes in SIX1 level, we then performed Co-IP assay to confirm the interactive relationship. IF staining further displayed that SPOCK1 and SIX1 were complexed together in cytoplasm. Li *et al*. put forward that SIX1 participated in the transcriptional regulation of the Warburg effect in BC (19), providing a fatal evidence for SIX1 to function as a hallmark of cancer. Herein, to further identified the special role of SIX1 in SPOCK1-mediated BC evolution, we blocked the expression of SIX1 using siRNA and explored the effect on cell cycle, proliferation, motility and EMT process. As we suspected, siSIX1 significantly abolished SPOCK1-induced facilitation of BC progression. Merely, a large of normative studies are still needed to clear this molecular mechanism.

In summary, this study contributed to illuminate the molecular mechanism by which SPOCK1 overexpression in human BC potentiated tumor progression. Our findings indicated that SPOCK1 is aberrantly overexpressed in BC. Interacting with SIX1 stimulated the AKT/mTOR signaling pathway to accelerate cell cycle progress, promote cell proliferation, triggered EMT progress and facilitated metastasis in BC (Fig. 7). SPOCK1 along with SIX1 might be prognostic factors for BC patients and promising therapeutic targets to prevent BC progression.

**Figure 7:**
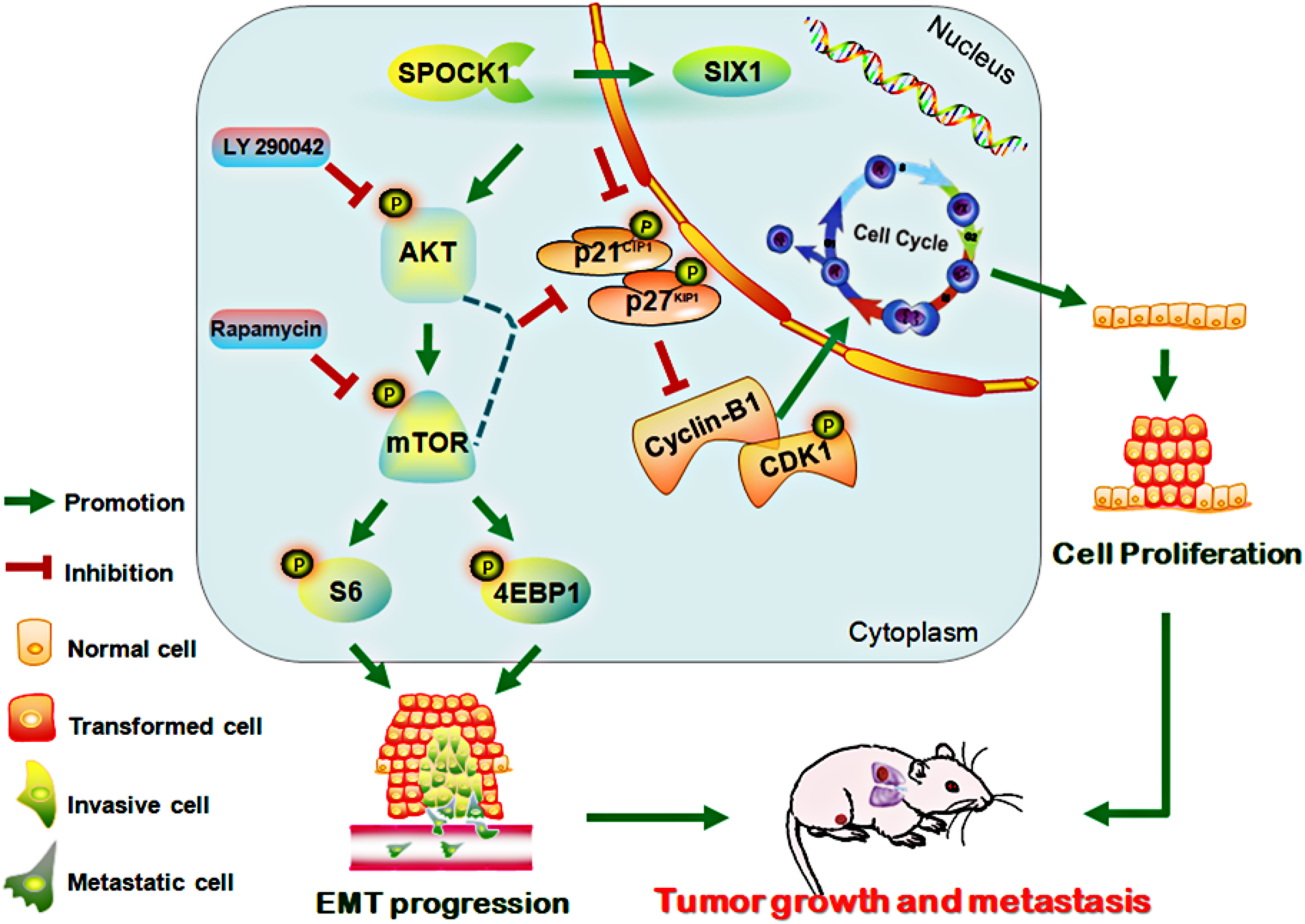
Schematic of proposed molecular mechanism of SPOCK1/SIX1-induced BC cancer cell growth and metastasis.

## Materials and methods

### Ethics statement

Our study complied the Declaration of Helsinki and approved by the Human Ethics Committee and the Research Ethics Committees of Yanbian University in China. All patients signed informed consent, which included consent to use the resection specimens for scientific research. All the specimens were kept in our tissue specimen bank and we promised to provide privacy protection for patients.

Animal care and experimental procedures in this study were approved by the Institutional Animal Care and Use Committee (IACUC) of Changchun Weishi testing technology service co. LTD and performed in accordance with the institutional guidelines.

### SPOCK1/SIX1 expression pattern

We performed SPOCK1/SIX1 mRNA expression in different cancers and confirmed the expression pattern of SPOCK1/SIX1 in BC by Oncomine database (https://www.oncomine.org/resource/login.html).

The comparison of different SPOCK1/SIX1 expression in various normal and cancer tissues and the expression of SIX1 protein in BC were used The Human Protein Atlas (HPA) (https://www.proteinatlas.org/) (22) and UALCAN (http://ualcan.path.uab.edu/analysis.html) (23) databases.

### Survival analysis

SPOCK1/SIX1 prognostic value in BC, including relapse free survival (RFS), distant metastasis free survival (DMFS), post progression survival (PPS) and overall survival (OS), was calculated by Kaplan-Meier plotter (http://kmplot.com/analysis/index.php?p=service&cancer=breast). Risk assessment was further assessed by SurvExpress (http://bioinformatica.mty.itesm.mx:8080/Biomatec/SurvivaX.jsp).

### Bioinformatics analysis

The protein and protein interaction networks in SPOCK1/SIX1 were established on the platform of GeneMANIA (http://genemania.org/) and Search Tool for the Retrieval of Interacting Genes (STRING) (https://string-db.org/cgi/input.pl) (24).

The heat map of the correlation between SPOCK1 and EMT markers in the same cohort was analyzed using UCSC Xena (http://xena.ucsc.edu/). The positive relationships for SPOCK1 and target genes were discerned and verified by Gene Expression Profiling Interactive Analysis 2 (GEPIA2) (http://gepia2.cancer-pku.cn/#index) (25).

### Cell culture

Human BC cell lines MCF7, MDA-MB-231, MDA-MB-453, MDA-MB-468, HS 578T, SKBR3 and normal immortalized mammary gland cell line MCF10A were cultured in Dulbecco Modified Essential Medium (DMEM) (Gibco, USA) with 10% FBS and 100 units penicillin and 100 mg/mL streptomycin.

### Plasmid construction and transfection

Human Lenti-shSPOCK1-GFP, Lenti-SPOCK1-GFP and negative control (Lenti-shNC and Lentivector control) were designed and packaged by Genechem (Co. Ltd., Shanghai, China). The target sequences of Lenti-shSPOCK1 were shown: 5’-TTTCGAGACGATGATTATT-3’ for shSPOCK1#2 and 5’-GCTGGATGACCTAGAATAT-3’ for shSPOCK1#3. The sequence of negative control was 5’-TTCTCCGAACGTGTCACGT-3’.

2×10^4^ BC cells were inoculated into 24-well plates for stable infection, then produce stably transfected cells by puromycin (2 μg/mL) after 48 h infection. The infection efficiency was identified by GFP gene reporter and western blot.

### Cell proliferation assay

The 3-(4,5-dimethyl-2-thiazolyl)-2,5-diphenyl-2-H-tetrazolium bromide (MTT) assay was used to detect BC cell proliferation. Briefly, infected BC cells were seeded in plates and grown to 80% confluences for staining with MTT (0.5 mg/mL; 100 μL; Dako, Denmark) and dissolved the crystal by dimethylsulfoxide per 24 h for 5 d. Three wells per group at least were analyzed and repeated three times.

### Colony formation assay

BC cells were inoculated in 6-well plates for incubating about 15 d. Then fixed the cells by methanol after being washed with cold PBS (Phosphate-buffered saline solution, Boster, China) and staining with Giemsa. Counting the colonies directly.

### 5-ethynyl-2’-deoxyuridine (EdU) incorporation assay

5×10^3^ BC cells seeded and grown in 96-well plates overnight and 100 μL 50 μM EdU medium (RiboBio, Guangzhou, China) per well were cultured for 2 h. Then the cells were fixed by methanol for 30 min and washing with PBS for 5 min twice. After permeabilizing with 0.5% TritonX-100 for 10 min twice and washing with PBS for 5 min, 1×Apollo dye was used to stain the cells for 30 min, repeated washing. Finally, the signal was visualized and recorded by a microscope after Hoechst 33342 counterstaining.

### Flow cytometry assay

Washing the cells with 10 mL cold PBS, then centrifuging at 1000 rpm for 10 minutes and discarding the supernatant. Added 5 mL of 75% ethanol and incubated at −20 °C overnight. Next day, washed twice with cold PBS to remove the ethanol, and centrifuged the cells for 10 minutes at 1500 rpm and discarded supernatant. Resuspension the cell pellets in 0.5 mL of PI/RNase Staining Buffer. After incubating 15 min at room temperature, stored tubes on ice freed from light prior to analyzing. Finally, samples were analyzed on the flow cytometer (BD Accuri C6) and used Modfit LT4.1 Software (Verity Software House, Inc., Topsham, ME, USA) to record the cell cycle distribution.

### Immunofluorescence (IF)

BC cells were attached with 4% paraformaldehyde. Permeabilizing cells with 0.5% TritonX-100 for 15 min and blocking with 3% Albumin Bovine V (Solarbio, Beijing, China) for 2 h. Then incubating primary antibodies overnight at 4 ℃. Incubating with second antibodies (A1108, Invitrogen, USA), and counterstaining by DAPI with an Antifade Mounting Medium (Beyotime, Shanghai, China) the next day. Finally, cover-slips signal was captured by the microscope.

### Wound healing assay

BC cells were seeded and grown in 6-well plates at approximately 80% confluences, scratching the monolayer uniformly on the surface by 200 μL. The scratched areas were recorded by microscope at 0h, 24h, 48h, and 72h. The migration distances were measured by Image J software for analyzing.

### Migration and invasion assay

3×10^4^ BC cells were inoculated onto the upper chambers with or without coating Matrigel (BD Biosciences) containing 1% with serum in DMEM. Filling 20% fetal bovine serum/DMEM media into the lower chamber. The chambers were attached with 4% paraformaldehyde for 5 min after incubation for 24-48 h at 37 °C, 5% CO_2_. Then staining with Giemsa after washing with cold PBS. The migrated cells were counted by a microscope. The experiment was performed three times to reduce the possible effects of biological variability.

### siRNA transfection

The sequence of si-SIX1 was SIX1-siRNA: 5’-GGGAGAACACCGAAAACAA-3’. BC cells were transfected with SIX1 siRNAs or control siRNA using Lipofectamine™ 3000 Reagent (Invitrogen, USA) according to the manufacturer’s instruction.

### Western blot

BSA Protein Assay Kit (A8020-5, Roche, Basel, Switzerland) was applied to measure the protein concentration after lysing BC cells with RIPA buffer. The total protein was dissolved by SDS-PAGE loading buffer and transferred onto poly vinylidene fluoride (PVDF) membranes (Millipore, Billerica, MA, USA). The primary antibodies (Table 3) were incubated overnight at 4 °C. Second day, the HRP-conjugated secondary antibodies (CST, Danvers, MA, USA) were incubating for an hour, then antibody-reactive bands were visualized via enhanced chemiluminescence (ECL) system (Millipore, USA) in an Imager. Operations were as follows (Fig.8):

**Table 3.** The information of antibodies

| Antibodies | Source company | Dilution ration |
| --- | --- | --- |
| Anti-SPOCK1 | R&D Systems, USA | 1:1000 |
| Anti-GAPDH | Proteintech, Wuhan, China | 1:1000 |
| Anti-p21(CDNK1A) | Proteintech, Wuhan, China | 1:1000 |
| Anti-p27[Kip1] | Proteintech, Wuhan, China | 1:1000 |
| Anti-CDK1 | Proteintech, Wuhan, China | 1:1000 |
| Anti-Cyclin-B1 | Cell signaling Technology, USA | 1:1000 |
| Anti-Survivin | Cell signaling Technology, USA | 1:1000 |
| Anti-C-myc | Santa Cruz, Dallas, Texas, USA | 1:1000 |
| Anti-Slug | Elibscience, Wuhan, China | 1:1000 |
| Anti-E-cadherin | Elibscience, Wuhan, China | 1:10000 |
| Anti-Vimentin | Cell signaling Technology, USA | 1:1000 |
| Anti-Snail | Cell signaling Technology, USA | 1:1000 |
| Anti-ZO-1 | Cell signaling Technology, USA | 1:1000 |
| Anti-Twist | Abcam, USA | 1:500 |
| Anti-S100A4 | Abcam, USA | 1:1000 |
| Anti-ZEB1 | Proteintech, Wuhan, China | 1:1000 |
| Anti-SIX1 | Sigma, USA | 1:1000 |
| Anti-p-AKT | Cell signaling Technology, USA | 1:2000 |
| Anti-AKT | Cell signaling Technology, USA | 1:1000 |
| Anti-p-mTOR | Cell signaling Technology, USA | 1:1000 |
| Anti-mTOR | Cell signaling Technology, USA | 1:1000 |
| Anti-p-S6 | Cell signaling Technology, USA | 1:1000 |
| Anti-S6 | Cell signaling Technology, USA | 1:1000 |
| Anti-p-4EBP1 | Cell signaling Technology, USA | 1:1000 |
| Anti-4EBP1 | Cell signaling Technology, USA | 1:1000 |

**Fig 8.**
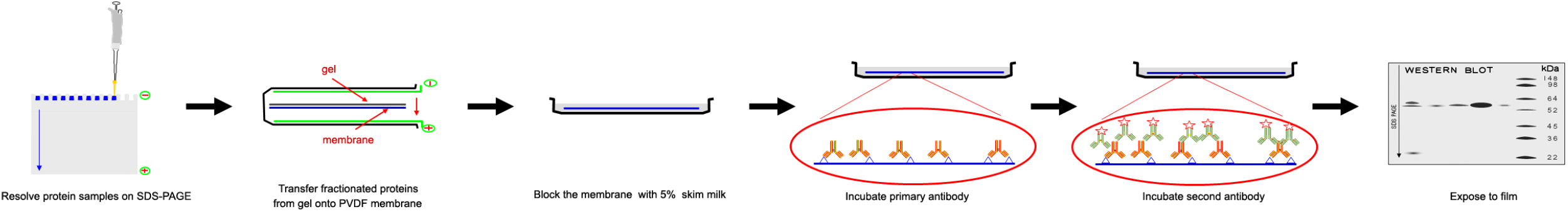
Schematic diagram of Western blot operations.

### Co-Immunoprecipitation (Co-IP)

In brief, added lysis buffer and incubated on ice for 30 min after washing the cell pellet with cold PBS. Scraped cells and centrifuged about 5 min, 15000 rpm, then transfer supernatant to a new microcentrifuge. Preclear the Protein A/G PLUS-Agarose beads (Santa Cruz, USA) with cold PBS and pre-blocked with BSA (Bovine serum albumin fraction V, Solarbio, China) to reduce non-specific immunoglobulin binding. Pellet beads, control IgG and 200 μL cell lysate incubated at 4 °C about 1 h, transfer supernatant to a fresh microcentrifuge tube on ice and added 5 μL primary antibody and incubate overnight at 4 °C, after centrifugation at 6500 rpm for 1 min at 4 °C. Cap tubes and incubated at 4 °C on a rocker platform overnight. Collected immunoprecipitates by centrifugation at 2500 rpm for 1 min at 4 °C. Aspirated and discarded supernatant carefully. Washed pellet 3 times with lysis buffer, each time repeating centrifugation step above. After final wash, carefully aspirated and discarded supernatant and resuspended pellet in 20 μL of 3× electrophoresis sample buffer. Boiled samples for 5 minutes and analyze 25 μL aliquots by western blot.

### Mouse xenograft model

To establish the orthotopic BC model, MDA-MB-231 cells stably overexpressing SPOCK1 (MDA-MB-231-SPOCK1) and MCF7 and SKBR3 cells stably silenced SPOCK1 (MCF7-shSPOCK1; SKBR3-shSPOCK1) as well as their negative control (MDA-MB-231-Vector; MCF7-NC; SKBR3-NC) were implanted in the mammary gland fat pad of BALB/c nude female mice (Viatal Rivers, Beijing, China). Injecting 10^6^ cells into the tail vein of 5-week-old nude mice for vivo lung metastasis models. Tumor sizes were monitored per 5 days, and volumes were calculated with a formula: Volume (mm^3^) = 0.5 × length × width ^2^. About 5 weeks, all mice were sacrifices, then tumors and lungs were removed. The number of lung metastases was counted on the surface of the lungs. Finally, dissected tumors and lungs were hematoxylin and eosin staining.

### IHC staining analysis

In brief, tissue sections were deparaffinized, rehydrated and incubated with 3% H_2_O_2_ for 15 min. Then performing in sodium citrate buffer (pH 6.0) at 95 °C for antigen retrieval. After returning to the room temperature, the slides were incubated with primary antibodies overnight. Next day, secondary antibody was incubated for 2 h. The slides were developed in the reaction with a 3, 3’-diaminobenzidine chromogen and counterstained with Mayer’s hematoxylin. Positive control and isotope control selected the tonsil and Rabbit IgG, respectively. Negative control treated positive tissue sections with PBS instead of primary antibody. Immunostaining for SPOCK1 was judged by a double semi-quantitative scoring system (Table 4). Specific reference to our previous research (26).

**Table 4.** Immunohischemical scoring according to immune-staining intensity and area

| Staining area(%)<br>Staining intensity | "0"<br>(0-5%) | "1"<br>(5%-25%) | "2"<br>(25%-50%) | "3"<br>(50%-75%) | "4"<br>(75%-100%) |
| --- | --- | --- | --- | --- | --- |
| "0"(Negative) | Score0 | Score0 | Score0 | Score0 | Score0 |
| "1"(Weak) | Score0 | Score1 | Score2 | Score3 | Score4 |
| "2"(Moderate) | Score0 | Score2 | Score4 | Score6 | Score8 |
| "3"(Strong) | Score0 | Score3 | Score6 | Score9 | Score12 |

### Statistical analysis

Statistical analyses were carried out by SPSS version 17.0 software and Prism 8.0 for Windows. Chi-square tests (χ2) was used to compare the correlations between SPOCK1 expression and clinicopathological parameters. All data were displayed by mean ± standard deviation, which calculated for thrice experiments. One-way Anova was used to compare data between multiple groups, and pairwise comparisons between groups were performed by t-test. We considered *P*<0.05 as statistically significant.

## AUTHOR CONTRIBUTIONS

Z. L. conceived this study and takes responsibility for the quality of the data. M. X., X. W. and Y. Y. participated in the experiments. S. Z. and Z. L. played an important role in interpreting the results. M. X. and J. P. performed the data analysis. M. X. wrote the manuscript and X. Z. helped to modify the manuscript. All authors read and approved the final manuscript. The authors declare no conflict of interest.

